# Wheat shovelomics II: Revealing relationships between root crown traits and crop growth

**DOI:** 10.1101/280917

**Authors:** Shaunagh Slack, Larry M. York, Yadgar Roghazai, Jonathan Lynch, Malcolm Bennett, John Foulkes

## Abstract

Optimization of root system architecture represents an important goal in wheat breeding. Adopting new field methods for root phenotyping is key to delivering this goal. A novel ‘shovelomics’ method was applied for phenotyping root crown traits to characterize the Savannah x Rialto doubled-haploid (DH) population in two field experiments under irrigated and rain-fed conditions. Trait validation was carried out through soil coring on a subset of 14 DH lines and the two parents. We observed that drought reduced grain yield per plant by 21.0%. Under rain-fed conditions, nodal root angle and roots shoot^-1^ were positively associated with root length density (RLD) at 40-60 cm depth; RLD was also positively correlated with grain yield. Nodal root angle and roots shoot^-1^ were also positively associated with canopy stay green and grain yield under rain-fed conditions. We conclude that shovelomics is a valuable technique for quantifying genetic variation in nodal root traits in wheat, revealing nodal root angle and root number per shoot provide useful selection criteria in breeding programs aimed at improving drought tolerance in wheat.

**Highlight:** Nodal root angle and number shoot^-1^ measured using ‘shovelomics’ were positively associated with root density at depth and yield under drought in a Savanah x Rialto wheat DH population.

## Introduction

Wheat (*Triticum aestivum* L.) provides, on average, one-fifth of the total calorific input of the world’s population. In the UK, winter wheat is the most widely grown arable crop and contributes ca. 16 million tonnes per annum with an average productivity of c. 8.5 t ha^1^ (DEFRA, 2016). The significantly warmer and more extreme conditions now arising due to climate change (IPCC, 2014) mean that new cultivars with greater drought resistance must be developed to maintain food security. Worldwide, drought limits agricultural productivity more than any other single factor (Cattevelli et al., 2008). In the UK, water deficit limits wheat grain yields in some years, where the onset of drought is post-anthesis and losses are ca. 20-30% (Foulkes *et al*., 2002, 2016).

Wheat root systems consist of seminal roots that arise from primordia in the embryo and nodal roots, also referred to as crown roots, which originate from basal nodes of the main shoot and tillers. A wheat plant typically produces about 6 seminal axes and 10-15 crown root axes (Gregory et al., 1978). Extension of both seminal and nodal roots usually continues to flowering (Belford *et al.*, 1987; Gregory *et al.*, 2005) and the two root systems thus function in a complementary manner. The ability of a root system to acquire water is principally related to its root system architecture (RSA) affecting the root length density (RLD; root length per unit soil volume) and distribution with soil depth as well as the maximum root depth (van Noordwijk, 1983; Foulkes *et al.*, 2009; Wasson *et al.*, 2012). Deeper roots are generally required to enhance water uptake (Wasson *et al.*, 2012; Lynch, 2013; Richard *et al.*, 2015). Genetic variation in RLD at depth is reported in UK winter wheat cultivars in the field in the range 1-2 cm cm^-3^ (Ford *et al*., 2006; White *et al.* (2015) and the depth below which RLD is < 1 cm cm^-3^ was shallower in modern UK winter wheat cultivars at 0.36 m than in older cultivars released in the 1970s and 1980s at 0.86 m (White *et al.*, 2015). Improved grain yield under drought has been associated with increased root DW at depth in synthetic wheat derivatives in NW Mexico (Reynolds *et al.*, 2007).

A steeper angle and higher number of seminal roots in wheat seedlings have been linked to a more compact root system with greater density of roots at depth in wheat in Australia (Manschadi *et al.*, 2008, 2010; Olivares-Villegas *et al.*, 2007). The root angle of Japanese winter wheat cultivars, grown in controlled environments, was correlated with their vertical root distribution in the field (Oyanagi and Nakamoto, 1993). Studies in maize have also suggested that a steeper root angle is related to increased rooting depth under low nitrogen field environments in the USA and South Africa (Trachsel *et al.*, 2013). Steeper root angles have also been shown to be important under drought conditions in rice (Uga *et al*., 2013) and maize (Lynch, 2013).

The lack of high-throughput field phenotyping methods for root traits remains a bottleneck in breeding programs (Fiorani and Schurr, 2013). Field phenotyping methods for roots such as rhizotrons, mini-rhizotrons and assessments of root parameters from soil cores (root washing and image analysis) are generally low to medium throughput. Higher throughput field phenotyping techniques include the soil-core break method (Köpke, 1979) and ‘shovelomics’ (Trachsel *et al.*, 2011). In the core-break method, soil-root cores extracted from the field are broken transversely and the roots on the exposed cross sections counted (Manske, 2001). The number of roots visible is then used to estimate RLD from established calibrations. A field study in Australia on a range of genotypes (cultivars, near-isogenic lines and recombinant inbred lines) showed this technique was able to identify directly variation in deep root traits (Wasson *et al*., 2014). However, this method works best on clay soils and is not suitable for all soil types. Shovelomics (or root crown phenotyping) involves the excavation and visual scoring of root crowns extracted from the field. Results in maize have been shown to be well correlated with root depth and root system total length (Trachsel et al., 2011). Shovelomics has been shown to be a useful tool for quantifying genetic variation in maize (Trachsel *et al*., 2011; Lynch, 2011; Abiven *et al.*, 2015), barley (Wojciechowski *et al.*, 2015) and durum wheat (Maccaferri *et al*., 2016). We have developed a high-throughput shovelomics technique for phenotyping root crown architecture of the whole root crown as well as the main shoots and tillers in bread wheat (see co-submitted paper by York et al., 2018).

The present study reports associations between nodal root traits measured using the new bread wheat shovelomics technique in irrigated and rain-fed field conditions in two years. We validate our results using soil coring in a subset of Savannah x Rialto DH population lines. Our results reveal that nodal root angle and root number per shoot represent valuable selection criteria in wheat breeding programs to improve drought tolerance.

## Materials and methods

### Experimental design, treatments and plot management

Two field experiments were carried out in 2013-14 and 2014-15 (referred to hereafter as 2014, 2015, respectively) at the University of Nottingham farm, Leicestershire, UK (52.5°N, 1.3°W). Each experiment used a randomised block, split–plot design, in which two irrigation treatments (fully irrigated and unirrigated) were randomised on main-plots, and 94 Savannah x Rialto doubled-haploid (DH) lines and the two parents were randomised on sub-plots (1.65 x 6.0 m) in two replicates. Both parents are semi-dwarf (*Rht-D1b*) UK winter wheat cultivars. Rialto was bred is suitable for some bread-making processes, and was released by RAGT Seeds Ltd in 1995. Savannah is a feed wheat cultivar, and was released by Limagrain UK Ltd in 1998. The soil was a sandy medium loam to 80 cm over Kyper marl clay of the Dunnington Heath series. In the irrigated treatment, a trickle irrigation system was used to maintain soil moisture deficit, calculated using the ADAS Irriguide model (Bailey & Spackman, 1996), to < 0.50 available water (AW) up to GS61 + 28 days and <0.75 AW thereafter. The AW capacity to 1.2 m soil depth was 176 mm. No water was applied in the unirrigated treatment.

Previous cropping was spring oilseed rape in 2013 and winter oilseed rape in 2014. The experiments were sown on 19 November 2013 and 20 October 2014. In each experiment, the field was ploughed and power harrowed and rolled after drilling. Seed rate was adjusted by genotype according to 1,000 grain weight to achieve a target seed rate of 320 seeds m^-2^; rows were 0.13 m apart. In each season, 200 kg ha^-1^nitrogen fertilizer as ammonium nitrate was applied in a three-split programme. P and K fertilizers were applied to ensure that these nutrients were not limiting. Plant growth regulator was applied at GS31 to reduce the risk of lodging. Herbicides, fungicides and pesticides were applied as required to minimise effects of weeds, diseases and pests.

### Crop measurements

A subset of 14 DH lines (Lines 1, 17, 19, 29, 32, 37,40, 44, 47, 64, 72, 73, 88, 98) and the two parents was selected for additional sampling by soil coring for assessment of RLD in 2014 and 2015. All other crop measurements were carried out on all 94 DH lines and the two parents in 2014 and 2015.

### Crop developmental stages and plant establishment

Dates of GS61 (anthesis) and GS89 (physiological maturity) were recorded according to the Zadoks growth stage (Zadoks *et al.* 1974) in all sub-plots in each year. The growth stage for the sub-plot was taken as when more than 50 % of the shoots were at the specific stage. Physiological maturity (GS89) was assessed as the date when less than 20% of the stem area was remaining green. In 2014-5, images of each sub-plot were taken after emergence on 10 November 2014 using a digital camera (Canon EOS 700D/T5i DSLR) and then processed in Image J to estimate a sub-plot percentage ground coverage. The percentage ground coverage was taken as an estimate of plant establishment.

### Shovelomics root crown assessment

The methodology for assessing root crown traits in 2015 was as described by York *et al.* (2018). The traits measured were nodal root number per plant, root number per shoot, root length and root angle. In 2014, the same methodology was used, except that the nodal root traits were visually scored using a protractor and ruler. Briefly, root crowns were excavated from all sub-plots during early to mid-grain filling on 25-26 June 2014 and on 8-9 July 8-9 2015. Three plants were selected from an internal row based on their uniform size in relation to neighbouring plants. A spade was inserted to 15 cm depth on either side of the focal plants with the width of the blade parallel to the row. The focal plants and attached soil were lifted from the ground and placed in a plastic bag and transported to a washing station where the roots with attached soil were placed into a 10 L bucket filled with water for 10 minutes to allow the soil to loosen (Fig. 1(a)). The root crowns were then sprayed with low pressure water from a hose to remove remaining soil. The number of fertile shoots (those with an ear) for each of the three plants was counted. The root crowns were then severed by cutting approximately 1 cm above the base, and stored at 4°C prior to assessment.

**Fig. 1.**
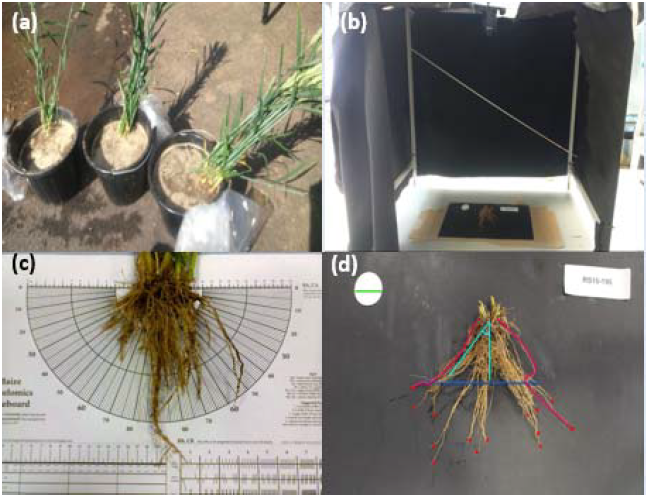
(a) Root crowns soaking in the field, (b) root crown imaging station in the laboratory, (c) root crown analysed using visual scoring board and (d) root crown analysed using ImageJ software.

In 2014, the root crowns were placed on a flat, white surface and the whole root crown and shoot base was visually scored for the following traits: shoot number, nodal root length, angle, and number of root axes. The number of nodal root axes was counted. Nodal root angle was measured for the outermost root axes on the left and right-hand sides of the crown, at 5 cm depth, using a protractor. Nodal root lengths were measured for the outermost roots on the left and right-hand sides of the root crown using a ruler (Fig. 1 (c)).

In 2015, shovelomics root traits were assessed as described by York *et al.* (2018) but in the current analysis only whole crown properties are used to keep consistency with the 2014 measurements. Root samples were excavated and washed as described above for 2014 and the whole root crown and shoot base was imaged using a digital camera attached to an aluminium frame covered in black cloth to minimize directional lighting and maximize diffuse lighting in the laboratory (Fig, 1(b)). The root crowns were placed on a matte black vinyl background with a 42 mm circular filter paper for scaling and a sample ID label. The camera was a Canon EOS 700D/T5i DSLR with manual settings for the shutter time duration and aperture to optimize the root contrast with background. The images were then analysed using a project for the Object J plugin for Image J (Schneider *et al*., 2012) created to allow the angles, numbers, and lengths of crown and seminal roots to be measured from the whole crown. A polyline was used to measure the crown lengths of the outermost roots and the seminal root length, and the angles were derived trigonometrically from the width of the root crown at a distance from the shoot base of approximately 5 cm, consistent with 2014. For root number, each root axis was manually annotated and the count recorded in an output file (Fig, 1(d)). The image analysis gave values for the number of pixels corresponding to root length and numbers. Using the 42 mm circular scale, these pixel values were then converted to the relevant units for each root measurement using a programme written in R software (R Core Team, 2012). Thus, the image-based measurements accomplished the same tasks as manual.

### Soil coring, root washing, scanning and WinRhizo image analysis

For the subset of 14 DH lines and the two parents in the unirrigated treatment, two soil cores (4.2 cm diam. × 60 cm depth) per sub-plot (one within and one between rows) were taken using a hydraulic soil corer (FF Bond Engineering Solutions, UK). The cores were divided into three 20 cm soil depth horizons (0-20, 20-40 and 40-60 cm) and the samples stored at 4 °C for up to 14 d prior to root extraction. The roots were extracted using a Gillison’s root washer (Gillison Variety Fabrication, Benzonia, MI). This system separates roots from soil using pressurized spray jets and low energy air flotation, causing the roots to float through the overflow pipe into a sieve. Each sample was left in the root washer for 10 minutes. The extracted roots were then stored in a freezer (−20 °C) in 50 ml plastic bottles prior to image analysis.

The clean root samples were imaged at 500 DPI resolution using an Epson Expression 11000XL scanner (Seiko Epson Corporation, Tokyo, Japan) with a transparency adapter. A preliminary study found a perfect correlation of derived root lengths from images scanned as JPEG or TIFF so scans were saved as JPEG to use efficiently storage. The scanned images were analysed using the WinRHIZO regular V.2002c software (Regent Instruments Inc., Quebec, Canada). After scanning each sample, the dry weight was recorded after drying for 48 h at 80¼.

The distribution of root length with soil depth was estimated according to equation (1) (Gale and Grigal, 1987):

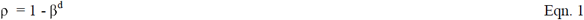

where ρ is the fraction of the root system accumulated from the soil surface to a given depth (d) and β is a parameter that describes the shape of the cumulative distribution with depth (Jackson *et al*., 1996).

### Grain yield and above-ground DM

In each experiment, approximately 50 shoots per sub-plot were hand-harvested by cutting at ground level at physiological maturity (GS89). In the laboratory, shoots were separated into fertile (those with an ear) and infertile shoots and counted. The fertile shoots were separated into ears and straw. A 25% sub-sample of the straw was taken (by fresh weight) and weighed. The dry weight of the ears and the sub-sample of the straw were recorded after drying at 80^°^C for 48 h. After threshing the ears using a Wintersteiger KG threshing machine (Wintersteiger, Austria), the dry weight of the grain, chaff and straw was weighed separately after drying for 48 h at 80°C. Five hundred grains from a sub-sample of grain (ca. 20 g) were counted by a Contador seed counter (Pfeuffer, Germany) and weighed to obtain the 1,000 grain weight. From these data the grain DM and grain DM per fertile shoot were calculated. The grain yield and above-ground DM per plant were then calculated by multiplying the grain yield and above-ground DM per fertile shoot by the fertile shoot number per plant assessed on the three plants per sub-plot used for the shovelomics assessments.

### Flag-leaf senescence and Normalised Difference Vegetation Index (NDVI)

Flag-leaf senescence was measured from anthesis (GS61) to full senescence every 3-4 days using a visual senescence score chart ranging from 0 - 10 (0; fully green and 10; fully senesced) as described by Gaju *et al*. (2011). Visual assessments were carried out for each sub-plot in both the irrigated and unirrigated treatments, and the values fitted against thermal time (post GS61; base temperature 0^°^C) applying a logistic regression equation using GenStat 18^th^ edition software package (Payne *et al.*, 2012; VSN International, Hemel Hempstead UK) as:

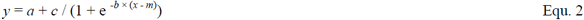

where *y* is the visual senescence score; *x* is thermal time from GS61 (base temperature 0°C); a is the lower asymptote; *m* is thermal time for point of inflection; *b* is the slope at the point of inflection; and *a*+*c* is the upper asymptote.

The onset of flag-leaf senescence (SEN_ONSET_) was taken as the thermal time when the flag-leaf senescence score was 2.0 and the end of flag-leaf senescence (SEN_END_) as the thermal time when the flag-leaf visual senescence score was 9.5. Senescence rate (SEN_RATE_; slope) was estimated as *b*. Values were calculated for each sub-plot and the fitted values were subjected to ANOVA.

The Normalized Difference Vegetative Index (NDVI) spectral reflectance index was measured using a handheld Greenseeker spectroradiometer (Trimble Navigation Ltd, USA). All sub-plots were measured on the same dates, at approximately GS61+35 days in each season (12 July 2014 and 13 July 2015). The Greenseeker spectroradiometer was held 50 cm above the crop canopy. A reading was taken per sub-plot when the sky was clear and there was sufficient radiation (Pask *et al.*, 2012) and NDVI was then calculated as in Equation 3 (Gutiérrez-Rodríguez *et al.*, 2004).

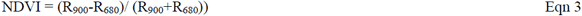

Where R_900_ is the reflectance in the near infrared band at 900 nm and R_680_ is the reflectance in the red visible band at 680 nm.

### Statistical analysis

For both years, GenStat 18^th^ edition (VSN International, Hemel Hempstead, UK) was used for statistical analysis of variance (ANOVA) of traits applying a split-plot design with replications regarded as random effects and genotypes as a fixed effects, and the least significant difference (LSD) test was used to compare the means between specific treatments. For ANOVAs across years, Bartlett’s test (P=0.05) was used to test for the homogeneity of variances, and years were regarded as random effects. The same software was used for correlation and linear regression analysis using the mean values for genotypes in the irrigated and unirrigated treatments. R version 3.4.3 was used to create the principal component analysis bi-plots (R Core Team, 2013).

## Results

### Drought effects on plant growth

Averaging across years, drought reduced grain yield plant^-1^ from 11.6 to 8.93 g (−21%) (p=0.06; Table 1). DH lines ranged from 7.03-18.3 g plant^-1^ under irrigated and 4.4-14.2 g plant^-1^ under unirrigated conditions (p=0.004). Yield decreases under drought ranged amongst lines from 0.51 to 9.51 g plant^-1^ (p< 0.001). The reduction in yield plant^-1^ was mainly associated with reduced above-ground DM (AGDM) plant^-1^ rather than harvest index. AGDM plant^-1^ decreased from 18.9 g under irrigated conditions to 14.6 g under drought (p< 0.001). Decreases under drought ranged amongst the DH lines from 0.10 - 13.96 g plant^-1^ (p<0.001).

**Table 1.** Maximum, minimum, mean, upper quartile and lower quartile values for GY_P_; Grain yield per plant, AGDM_P_; Above-ground dry matter plant^-1^, TGW; Thousand grain weight, PH; Plant height, SNp; Fertile shoot number per plant, CRA; Crown root angle, CRL; Crown root length plant^-1^, CRNp; Crown root number per plant and CRNs; Crown roots per shoot under irrigated and unirrigated conditions, and SED. Values represent means of 2014 and 2105.

| Treatments |  | Gy<sub>p</sub><br>g | AGDM<sub>p</sub><br>g | TGW<br>g | PH<br>cm | SN<sub>p</sub> | CRA<br>° | CRL<br>cm | CRN<sub>p</sub> | CRNs |
| --- | --- | --- | --- | --- | --- | --- | --- | --- | --- | --- |
| Irrigated | Max | 20.7 | 39.2 | 48.9 | 83.8 | 9.50 | 54.7 | 14.2 | 69.5 | 10.2 |
|  | Min | 5.87 | 9.86 | 34.8 | 59.9 | 3.00 | 44.5 | 6.55 | 24.8 | 6.02 |
|  | Mean | 10.4 | 17.2 | 42.1 | 72.3 | 4.75 | 48.9 | 10.1 | 37.3 | 8.18 |
|  | Upper Quartile | 11.7 | 19.0 | 43.8 | 75.9 | 5.25 | 52.3 | 11.8 | 40.1 | 8.68 |
|  | Lower Quartile | 8.73 | 14.8 | 40.2 | 69.2 | 4.00 | 44.5 | 8.18 | 32.3 | 7.69 |
|  | Rialto | 11.6 | 19.7 | 45.4 | 78.5 | 4.25 | 36.1 | 9.38 | 28.3 | 7.08 |
|  | Savannah | 15.8 | 25.4 | 47.1 | 77.7 | 6.00 | 36.4 | 11.7 | 39.0 | 6.79 |
| Unirrigated | Max | 20.7 | 31.9 | 53.5 | 83.6 | 8.50 | 64.7 | 12.0 | 69.5 | 10.2 |
|  | Min | 5.87 | 9.85 | 35.7 | 58.1 | 3.00 | 43.1 | 6.55 | 24.8 | 7.13 |
|  | Mean | 10.2 | 16.5 | 41.9 | 72.5 | 4.69 | 55.2 | 8.32 | 38.0 | 8.52 |
|  | Upper Quartile | 11.3 | 18.2 | 43.5 | 76.4 | 5.25 | 56.6 | 8.83 | 40.3 | 8.95 |
|  | Lower Quartile | 8.33 | 13.7 | 40.3 | 69.4 | 4.00 | 51.3 | 7.70 | 33.5 | 8.04 |
|  | Rialto | 9.12 | 14.7 | 44.9 | 73.7 | 4.75 | 52.6 | 8.99 | 37.5 | 8.11 |
|  | Savannah | 7.13 | 11.5 | 44.2 | 74.3 | 5.50 | 53.3 | 12.0 | 39.5 | 7.92 |
| SED (df) | Irrigation (1) | 0.23* | 0.39*** | 0.18** | 0.299** | 0.02*** | 1.89* | 0.21*** | 0.09** | 0.31*** |
|  | Genotype (95) | 1.14*** | 1.92*** | 1.72*** | 2.84*** | 0.33*** | 1.98*** | 0.48** | 1.62** | 0.88*** |
|  | Irr. x Gen. (95) | 1.62*** | 2.73*** | 2.42<sup>NS</sup> | 4.01<sup>NS</sup> | 0.46*** | 3.37* | 0.71* | 2.29** | 1.24*** |
|  | Yr x Irr. x Gen. (95) | 1.93<sup>NS</sup> | 3.31<sup>NS</sup> | 2.80<sup>NS</sup> | 6.11<sup>NS</sup> | 0.64<sup>NS</sup> | 4.07<sup>NS</sup> | 0.95<sup>NS</sup> | 3.18* | 1.72*** |

: \*\*\*denotes p<0.001; \*\*p<0.01 and \*p<0.05 significance levels; ns = not significant

There was no significant effect of drought on 1,000 grain weight (p=0.71). DH lines overall ranged from 33.3-50.9 g (p<0.001). There was no irrigation × genotype interaction. Overall plant height decreased slightly from 69.4 cm under irrigated to 67.8 cm under rain-fed conditions (p<0.001). The DH lines ranged from 56.1 to 88.7 cm (p<0.001); there was no irrigation × genotype effect. For grain yield and yield components the year x irrigation x genotype interaction was not statistically significant.

Drought reduced fertile shoots plant^-1^ from 5.3 to 4.3 (−34%) (p=0.007, Table 2). Genotypes ranged from 3.3-10.3 shoots plant^-1^ under irrigated conditions, and 2.3-7.9 shoots plant^-1^ under rain-fed conditions (p<0.001). Decreases in shoot number under drought ranged amongst DH lines from 0.6-54.9% (p<0.001) and there was a year × irrigation × genotype interaction (p<0.001).

**Table 2.**
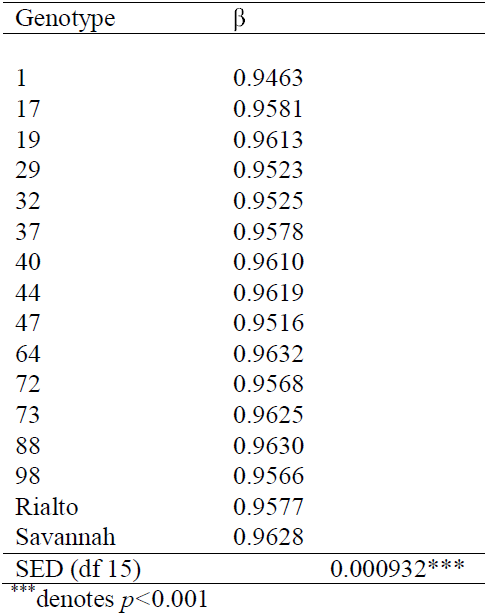
β value for relative root length density distribution (0 - 60 cm soil depth) with depth at harvest for 14 Savannah x Rialto DH lines and the Rialto and Savannah parents, Values represent means in 2014 and 2015 in unirrigated treatment.

| Genotype | $\beta$ |
| --- | --- |
| 1 | 0.9463 |
| 17 | 0.9581 |
| 19 | 0.9613 |
| 29 | 0.9523 |
| 32 | 0.9525 |
| 37 | 0.9578 |
| 40 | 0.9610 |
| 44 | 0.9619 |
| 47 | 0.9516 |
| 64 | 0.9632 |
| 72 | 0.9568 |
| 73 | 0.9625 |
| 88 | 0.9630 |
| 98 | 0.9566 |
| Rialto | 0.9577 |
| Savannah | 0.9628 |
| SED (df 15) | 0.000932*** |

\*\*\* denotes $p < 0.001$

### Shovelomic profiling of root crown traits

Averaging across years, drought reduced nodal roots plant-1 (NRNp) by 12.2 (−18.8%) (p=0.007; Table 1). DH lines ranged from 40.0-54.1 roots plant-1 under irrigated conditions and 20.7-54.1 roots plant-1 under drought (p<0.001). Decreases under drought ranged from 0.17% - 55.73% (p<0.001) and there was a year × irrigation × genotype interaction (p<0.001). Turning to consider average number of nodal roots per shoot (NRNs), drought increased NRNs from 7.9 to 8.4 (6.3%) (p<0.001). DH lines ranged from 6.0-10.1 roots shoot-1 under irrigated conditions and 8.4-9.8 roots shoot-1 under rainfed conditions (p<0.001). Increases under drought ranged from 0.3% - 22.8% (p<0.001) and there was a year x irrigation x genotype interaction (p<0.001).

Drought increased crown root angle (greater angle representing steeper roots) by 14.9° (26.4%) (p=0.045). DH lines ranged from 18.5-46.1° under irrigated and 42.3-72.7° under unirrigated conditions (p<0.001). The increase in root angle under drought ranged across genotypes from 5.68% - 45.6% (p<0.001) and there was no year x irrigation x genotype interaction (p<0.001). Nodal root length was reduced under drought overall by 6.5 cm (−46.0%) (p< 0.001). Genotypes ranged from 11.6-16.6 cm under irrigated conditions and 5.9-9.7 cm under rain-fed conditions (*p*<0.001). Decreases under drought ranged from 37.4-67.4% (p<0.001); there was no year x irrigation x genotype interaction.

There was a positive linear association between fertile shoots plant^-1^ and NRN_p_ under irrigated and unirrigated conditions (R^2^ 0.67, p<0.001 and R^2^ 0.68, p<0.001, respectively).

### Root length density at depth and relationship with shovelomic traits

Root length density (0-60 cm) in the rainfed treatment in the subset of 16 genotypes ranged from 1.60-1.97 cm cm^-3^ in 2014 (p=0.03) and 1.27-1.39 cm cm^-3^ (L 98) in 2015 (p=0.04; Fig 2). In the 40-60 cm soil layer, RLD ranged from 0.84-1.58 cm cm^-3^ in 2014 (p = 0.01) and 0.78-1.48 cm cm^-3^ in 2015 (p< 0.05). Overall different relative RLD distributions with depth were apparent amongst the genotypes with lines 44 and 64 showing higher β values (relatively deeper root distribution) than lines 1 and 47 (Table 2).

**Fig. 2.**
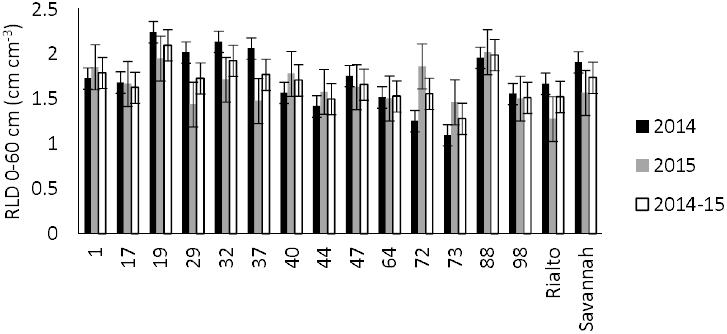
Root length density 0-60 cm for 14 Savannah x Rialto DH lines and 2 parents under rainfed conditions in 2014 and 2015 and mean 2014-5, error bars are standard errors of mean (SEM).

**Fig. 3.**
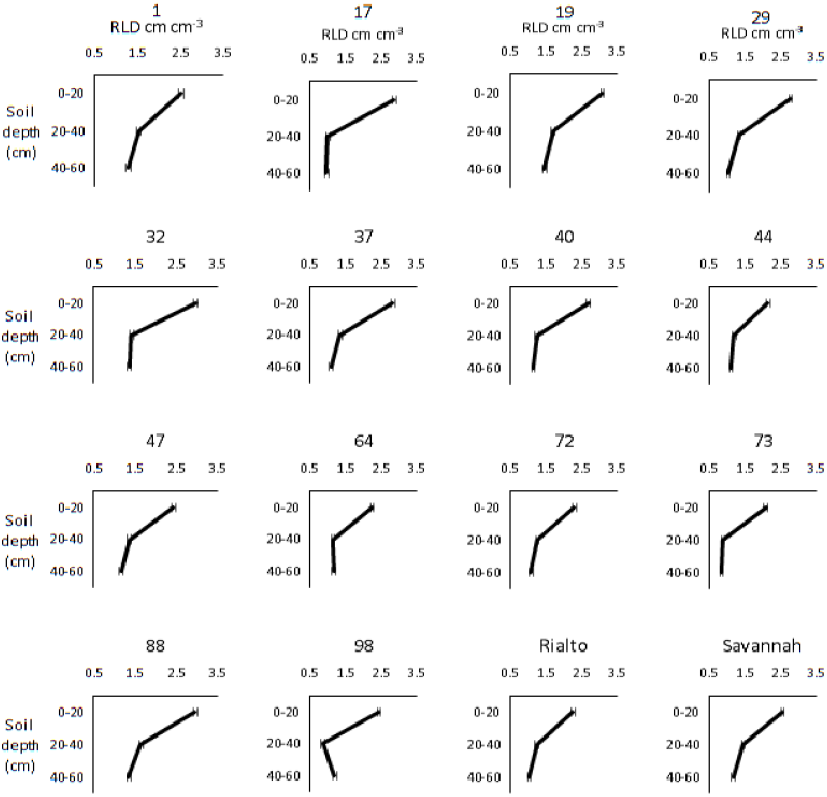
Root length density (RLD; cm cm^-3^) for 0-20, 20-40 and 40-60 cm soil layers for 14 Savannah x Rialto DH lines and two parents,, error bars are standard error of mean (SEM). Standard error of difference of mean (SED) for genotype (df 15) for 0-20 cm: 0.0718; 20-40 cm: 0.0950 and 40-60 cm layers: 0.1053. Values represent means in 2014 and 2015 in unirrigated treatment.

Averaging across years in the rainfed treatment, there was a positive linear relationship amongst the 16 genotypes between RLD at 40-60 cm and each of nodal root angle (R^2^ 0.51; p=0.04; Fig. 4a), nodal root number shoot^-1^ (R^2^ = 0.55, *p*< 0.03; Fig. 4c) and nodal root length (R^2^ 0.55, *p*< 0.03; Fig. 4e). There was also a negative linear association between RLD at 40-60 cm and crown root system width (R^2^ 0.49. *p*< 0.05; Fig. 4d). There was no association between nodal root number plant^-1^ and RLD at 40-60 cm. RLD at 40-60 cm was also positively associated with grain yield plant^-1^ in the rain-fed treatment (R^2^ 0.51, *p*< 0.05; Fig. 4f).

**Fig. 4.**
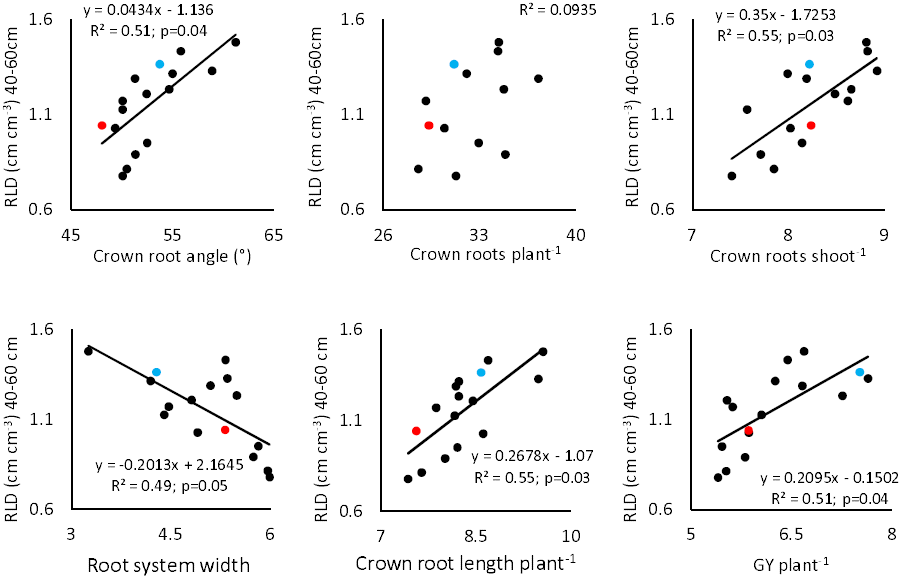
Relationship between root length density (RLD; cm cm^-3^) at 40-60 cm soil depth and (a) crown root angle, (b) crown roots plant^-1^, (c) crown roots shoot^-1^, (d) crown root system width, (e) crown root length plant^-1^ and (f) grain yield plant^-1^ for 14 Rialto x Savannah DH lines and two parents in irrigated and unirrigated treatment. Rialto and Savannah are indicated by red and blue circles, respectively. Values represent means in 2014 and 2015.

### Associations between shovelomic traits, stay-green and grain yield

Averaging across years in the rain-fed treatment, there was a positive linear relationship amongst the 94 DH lines between onset of flag-leaf senescence and grain yield plant (R^2^=0.23; p=0.04; Fig. 5a). RLD at 40-60 cm was also positively associated with onset of flag-leaf senescence in the subset of 16 genotypes (Fig. 5b). No significant association was found between onset of end of flag-leaf senescence and grain yield plant^-1^ under irrigated conditions.

**Fig. 5.**
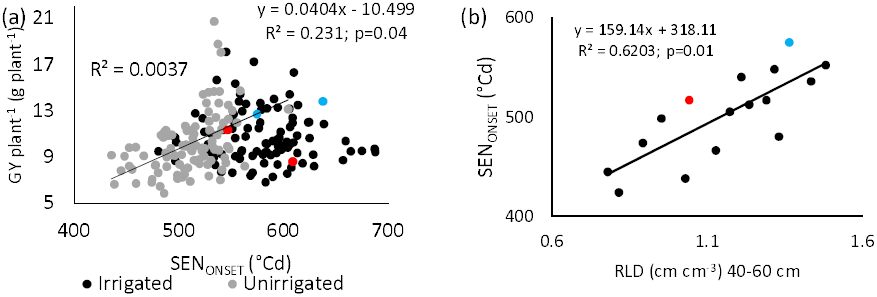
(a) Relationship between a) grain yield plant^-1^ and onset of flag-leaf senescence (SEN_ONSET_) for 94 Rialto x Savannah DH lines and the two parents under irrigated and unirrigated conditions and (b) SEN_ONSET_ and root length density at 40-60 cm for 14 Rialto x Savannah DH lines and the two parents in unirrigated treatment. Rialto and Savannah are indicated by red and blue circles, respectively. Values represent means across 2015 and 2016.

Biplots were created to examine the relationships amongst the root traits and above-ground traits in the irrigated and rain-fed treatments (Fig. 6). In the rain-fed treatment, the strong positive association between nodal roots shoot^-1^ and shoots plant^-1^ was confirmed, as well as the positive association between nodal root angle and each of grain yield plant^-1^ and AGDM plant^-1^. Nodal root angle was also positively associated with canopy stay-green (as indicated by NDVI at GS61+35d). In addition, TGW was positively associated with onset of flag-leaf senescence. In the irrigated treatment, nodal root angle and nodal root number shoot^-1^ were not associated with grain yield plant^-1^. However, CRN_p_ was associated with grain yield plant^-^1. There were no associations between nodal root traits and senescence-related traits under irrigated conditions.

**Fig. 6.**
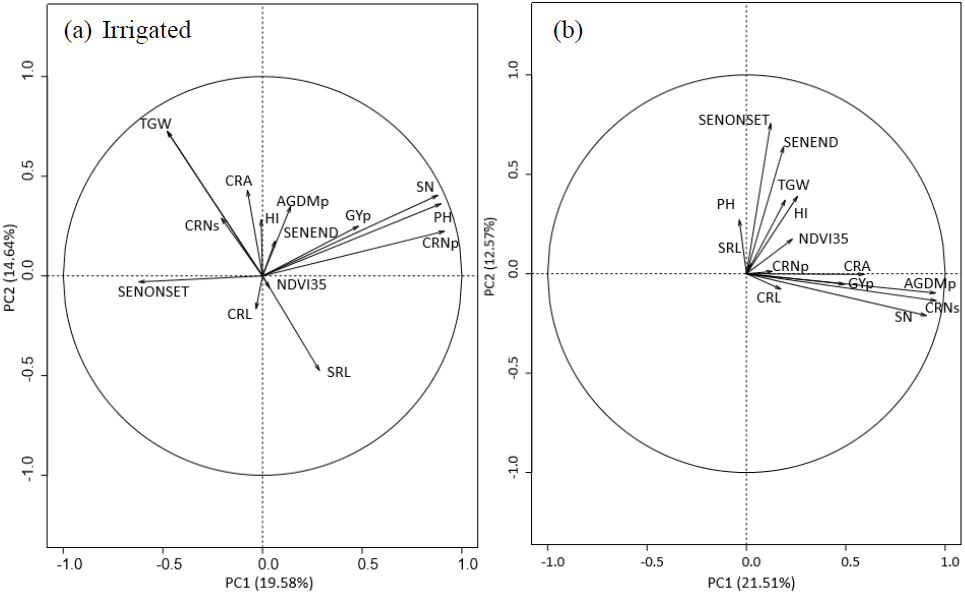
Principal component analysis of GYp; grain yield plant^-1^, HI; harvest index, TGW; thousand grain weight, AGDMp; above-ground dry matter plant^-1^, SN; fertile shoots plant^-1^; PH, plant height; CRA; crown root angle, SRL; seminal root length, CRL; crown root length, CRNs; crown roots shoot^-1^; CRNp; crown roots plant^-1^; NDVI35; NDVI at GS61+35 days; SEN_ONSET_, onset of flag-leaf senescence and SEN_END_; end of flag-leaf senescence) for 94 Rialto x Savannah DH lines and the two parents under (a) irrigated and (b) unirrigated conditions. Values represent means in 2014 and 2015.

## Discussion

The development of field-based high-throughput phenotyping for roots is a priority for drought research. Improved maintenance of yield under water stress has been demonstrated in wheat associated with deeper root systems (Sharma *et al.*, 2011; Ehdaie *et al.*, 2014).

### Shovelomic profiling of root traits and association with roots at depth

Shovelomics represents a high-throughput phenotyping method for field-grown crops and has been used to quantify genetic variation in root traits in maize (Trachsel *et al*., 2011; Lynch, 2011; Abiven *et al.*, 2015), legumes (Burrige *et al.*, 2016) and barley (Wojciechowski *et al*., 2015). Maccaferri *et al.* (2016) carried out field shovelomics for durum wheat recombinant inbred lines (RIL) for crown root length, number and angle and reported QTL. In our study we applied our shovelomics methodology for bread wheat (York et al., 2018) to quantify variation in nodal root angle, length, roots plant^-1^ and roots shoot^-1^ and association with RLD. The present range in nodal rot angle of 44.7-69.1° in the rainfed treatment was similar to that of 42.3-69.2° reported by Maccaferri *et al*. (2016) for the Colosseo × Lloyd durum wheat mapping population assessed at anthesis in the field under optimum agronomic conditions in Italy. However, values under irrigation of 31.7-48.7 ° were lower indicating that irrigation resulted in less steep roots as might be anticipated.

Our results showed a positive correlation between nodal root angle and RLD in the 40-60 cm soil layer in the rain-fed treatment. A more vertical angle of seminal roots of wheat seedlings was linked with more roots at depth in wheat in Australia (Manschadi *et al.*, 2008, 2010; Olivares-Villegas *et al.*, 2007), and root angle of Japanese winter wheat cultivars in controlled environments correlated with their vertical root distribution in the field (Oyanagi and Nakamoto, 1993). Previous studies in maize also found steeper root angle related to increased rooting depth under low nitrogen field environments in the USA and South Africa (Trachsel *et al.*, 2013). Modelling studies have also suggested that a steeper root angle in wheat may result in deeper roots and better maintenance of grain yield under drought (Manschadi *et al*., 2008). There was a negative relationship amongst the genotypes between nodal roots plant^-1^ and RLD in each soil layer, but a positive association with nodal roots shoot^-1^. The increase in the number of roots shoot^-1^ may have been due to the main shoot having many more roots than tillers and as tiller numbers reduced, the average number of roots per shoots weighted towards the main shoot. These results suggest that the wheat ideotype for deeper rooting may be a plant with relatively few tillers but a high number of nodal roots shoot^-1^. A field study in Pennsylvania in maize found that reduced nodal roots plant^-1^ led to increased root length at depth and 57% higher grain yield under water-stressed conditions (Gao and Lynch, 2016). In rice and wheat genotypes with fewer tillers were reported to have deeper (Yoshida & Hasegawa, 1982) or longer root systems (Duggan *et al*.,2005; Richards *et al*., 2006). The inverse relationship between tiller number and root density at depth could also partly relate to increased assimilate partitioning to the nodal roots associated with the main shoot and high order tillers which are the deepest roots having the longest residence times in the soil.

### Associations between rooting traits, stay-green and grain yield

In our experiments, there was a statistically significant, but mild drought with yield plant^-1^ reducing overall from 10.9 to 8.6 g plant^−1^ (−20.9%). This is representative of late-season drought effects reported for wheat in the UK with reductions of yield typically ca. 20-30% in dry years on drought-prone soil types (Foulkes *et al.*, 2001; 2002). In spite of the relatively mild drought stress, the range of yield reductions amongst cultivars was high as indicated by the significant irrigation x genotype interaction. Higher yield under irrigation was associated with greater yield loss under drought amongst the DH lines. From the physiological standpoint, it is not surprising that absolute reduction in yield for a given reduction in water resource is strongly influenced by yield potential (Fischer and Maurer, 1978; Foulkes *et al*., 2007; Aravinda-Kumar *et al*., 2011).

In the present study, increased RLD at 40-60 cm soil depth in rain-fed conditions was associated with delayed onset of flag-leaf senescence and higher grain yield and TGW. Thus, increased RLD at depth appeared to be a determinant of genetic variation in flag-leaf stay-green. Greater yield associated with longer green canopy area duration (stay-green) amongst genotypes has been reported under drought in wheat (Gorny and Garczynski, 2002; Verma et al., 2004; Foulkes et al., 2007; Christopher et al., 2008), sorghum (Borrell and Hammer, 2000) and maize (Campos et al., 2004). We found a positive correlation between onset of flag-leaf senescence and yield under drought, but no association under irrigation. The higher grain yield associated with stay-green under drought was likely due to source limitation of grain yield (Christopher et al., 2008; Bogard et al., 2011), and greener canopies have been reported to maintain the active photosynthetic rate better in wheat (Joshi et al., 2007).

In our study there was a large effect of drought on flag-leaf senescence timing; drought advanced onset of senescence by approximately 10 days. However, the grain yield decrease was relatively modest at 21%, suggesting that photosynthesis of non-laminar green organs (such as the ear, peduncle or sheaths) was still contributing to grain filling during the lamina senescence. Nevertheless, genetic variation in onset of flag-leaf senescence showed a moderately strong association with yield under the mild drought conditions (R^2^ 0.35; p=0.01). The mechanisms underlying the genetic differences in leaf senescence cannot be certain from present measurements. However, our results strongly imply that root traits were partly responsible for the stay-green effects with a positive association between flag-leaf senescence timing and RLD at depth (40-60 cm), which may represent a drought avoidance strategy that prohibits early onset of senescence due to drought. Under drought, stay green was previously associated with deeper roots under drought during the grain-filling period for two CIMMYT wheat lines SeriM82 and Hartog compared to check lines (Christopher *et al.*, 2008). It is important to note that RLD in this experiment was only measured to 60 cm soil depth and wheat roots have been recorded at depths of 1-2 m (White *et al*., 2015; Gregory et al., 1978), so present associations between nodal root traits and senescence traits and RLD require further validation and must be interpreted cautiously.

Our results showed genetic variation in the RLD distribution with depth according to β and positive relationships between β and nodal root angle (i.e. steeper root angle associated with relative deeper distribution of roots) (R^2^=0.23; p=0.04) and grain yield (R^2^=0.30; p=0.001) under drought. This is in agreement with previous studies showing improved yield under drought through distributing roots relatively deeper with soil depth in field conditions for bread wheat (Barraclough *et al*. 1989) and in soil columns in durum wheat (Carvalho *et al*., 2014). Using data for RLD distribution with depth from studies carried out on various field-grown winter wheat genotypes in the UK (Gregory *et al*., 1978; Barraclough, 1984; Barraclough & Leigh, 1984), King *et al*. (2003) calculated a range of β from 0.940-0.970 at harvest. This range was slightly higher than that obtained in our experiments in the range 0.9549-0.9637 in 2014 and 0.9573-0.9643 in 2015 under drought. Our results showed steeper nodal root angles were positively correlated with β under unirrigated conditions. Manschadi *et al*. (2008) also observed increased seminal root number and narrower root angles were associated with relatively deeper root distribution in Australia.

### Implications for plant breeding

Shovelomics is becoming an increasingly popular method for the high-throughput phenotyping of field-grown crop roots. The shovelomics method we have developed (York et al., 2018) and applied for phenotyping nodal root traits in winter wheat in the present study was shown to be a valuable technique. It was a relatively high-throughput method, making it possible to sample, wash and image root crowns from 384 plots in one person-week. Overall, analysis in ImageJ was preferable to visual assessment as it was significantly less time-consuming and was less dependent on the operator. This is significantly faster than the field soil coring, which took approximately one person-month to sample, wash and extract roots, and image the samples for 64 plots in the present study. The nodal root traits such as angle and number of roots per shoot had significantly positive relationships with RLD at 40-60 cm depth, which was positively associated with grain yield. However, the R2 values for these relationships were 0.5 - 0.6 and so up to 50% of phenotypic variation was not accounted for. Nevertheless, the present results demonstrated shovelomics can be used to measure nodal root traits which are an indirect indicator of root traits at 40-60 cm depth, potentially making it a useful field phenotyping technique; further work is required to determine whether root traits below 60 cm soil depth have equally strong relationships with nodal root traits, and the consistency of the correlations across a wider range of environments. Combining root system architectural properties with optimal rhizosphere qualities (York et al., 2016) will contribute further possible yield gains.

This high-throughput shovelomics platform could be used in future studies to phenotype large populations for root traits, such as nodal root angle, under drought stress to identify QTL, search for candidate genes and develop molecular marker for marker-assisted selection. There are examples of deploying QTL for root depth in other cereal species. In rice, the Dro1 gene related to steeper crown root angles and deeper rooting was identified by measuring nodal root traits in a high-throughput controlled environment study (Uga *et al*., 2011) and has since been used to produce drought tolerant NILs which have been phenotyped in field conditions (Uga *et al*., 2013).

## Acknowledgements

This work was funded by FP7-IDEAS-ERC FUTUREROOTS: Redesigning root architecture for improved crop performance ERC Future Roots Project ID: 294729. We thank Limagrain UK Ltd and John Innes Centre, UK for the use of the Savannah × Rialto DH population material.

